# A Bayesian method for estimation of plant soil water content with application to low-cost horticultural robotics

**DOI:** 10.64898/2026.07.14.738550

**Authors:** Alex J. Southgate

## Abstract

Climate change represents a challenge to food security by interfering with the environmental conditions needed for productive plant growth. While technology can be used for partial mitigation, access to technology is inequitable. Low-cost microcontrollers, such as the ESP32, have recently lowered the barrier for entry into prototyping smart devices. ESP32s equipped with capacitive moisture sensors have been suggested for low-cost smart plant watering systems. However, measuring moisture in soil is complex, potentially destructive, and requires careful calibration in order to characterise the response curve mapping soil water content to sensor measurements. Here, we developed a Bayesian method for estimating the inverse response curve from capacitive moisture sensor data, known water doses, and prior uncertainty, bypassing the need for destructive gravimetry. This method constitutes the core calibration module of the open-source *OpenHCult* software system for low-cost horticultural automation.

## 1 Introduction

Recent work has proposed the use of ESP32-class microcontrollers and low-cost moisture sensors for automated watering systems and measurement of soil water content (SWC) [2, 17, 20, 1], motivated by mitigating water shortages [1], with increased interest in application of IoT sensors to agriculture [15]. As global warming continues, both flooding and water shortages are increasing world-wide [4], including in the United Kingdom [12]. In previous work, ESP32/capacitive sensor calibration and validation has been performed by gravimetry [20]. However, it is expected that calibrated response curves depend on a range of factors, including soil type (i.e. clay content), and temperature [14]. Furthermore, end users may find it difficult to undertake calibration for each specific application type.

Typically, gravimetric sampling can be used to characterise SWC, but is destructive and time consuming; this process often involves drying soil by heating and therefore cannot be easily applied to existing plant systems. Capacitive moisture sensors, as with all electromagnetic methods (EM) measure SWC indirectly through the dielectric permittivity [9], though it is dependent on measurement electrical frequency [9, 18] and temperature [7]. Field capacity (FC), the water content at which internal drainage stops, is often utilized [8].

Ideal ranges for SWC varies across plant species and growth stages. For example, tomatoes (*Lycopersicon esculentum*) may grow best at 70% field capacity (FC) [13], while the cactus *Opuntia ficus-indica* was shown to exhibit stress under 30% FC [11].

Existing work has been undertaken on moisture sensor calibration and characterisation of variability. Bogena et al (2017) [3] developed a procedure for calibration of low-cost SMT1000 moisture sensors and undertook systematic calibration of hundreds of sensors in a range of media, observing sensor-to-sensor variability and both density and temperature dependence. In contrast, Kizito et al. (2008) [10] found that for research-grade ECH2O probes (Decagon Devices Inc., Pullman, WA) with a measurement frequency of 70 MHz, a single calibration curve was sufficient for relating electrical conductivity to SWC across different soil salinities, though they did find variability with the degree of probe immersion and dependency of measurements on temperature, and the sensors used are more an order of magnitude more costly than the capacitive sensors used in other studies (e.g. [20]). While cheap, these capacitive sensors can be reliable given controls on soil type and the ratio of solid matter to volume [16].

The objective of this work was to introduce an automated system and statistical method for automated calibration of sensors using watering volume data (dosage) rather than labour intensive gravimetric calibration. This aim was primarily motivated by the challenge end users may find with gravimetric calibration, as well as lack of generalisability of a given calibration curve.

## 2 Methods

### 2.1 Pre-processing

Irregularly spaced sensor data gathered from sensors was smoothed with an exponential moving average [6] (or exponentially weighted moving average (EWMA) [21]), a single-pole low-pass filter [19] with time-weighted coefficients:

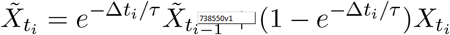

This was used to smooth high-frequency noise.

### 2.2 Calibration

#### 2.2.1 Integral data and the inverse response function

For sensor index *i* and time *t*, let *T*_*t*_ be the total water volume, *Y*_*i,t*_ the relative (here volumetric) SWC local to sensor *i*, and *X*_*i,t*_ = *h*(*Y*_*i,t*_) the sensor response. Let *V* be the container volume, and therefore *T*_*t*_*/V* the average volumetric SWC. *Y*_*t*_ and *T*_*t*_ cannot be measured directly non-destructively. If the inverse response function *h*^*™*1^ is known, *Y*_*i,t*_ = *h*^*™*1^(*X*_*i,t*_) could be computed. For multiple sensors, *T*_*t*_ and average volumetric SWC can be computed by averaging over *N* sensors, assuming sensors are uniformly distributed in the soil:

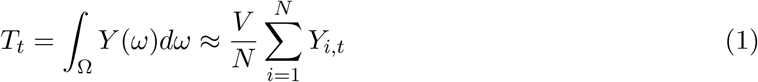

Since the response function can depend on both soil parameters, environmental factors, and sensor parameters including placement (e.g. how densely packed soil is around the sensor), we assume it is unknown and aim to estimate it.

Δ*T*, the change in total system water, is measurable for watering events that are recorded, which provides information on the response function. For example, if the inverse response was linear, assuming no noise, *T* = *V h*^*™*1^(*X*) could be reconstructed from (*X*_*i,start*_, *X*_*i,end*_, Δ*T* ), and the intercept from a single measurement *X*_*i*_ = *h*^*™*1^(0). In practice, the response curve is non-linear, and plants are usually sensitive to water content, tolerating only a narrow range of saturation, so the values (*X*_*i,start*_, *X*_*i,end*_, Δ*Y*_*i*_) are drawn from a narrow range of the domain. As such, only a narrow section of the curve *h*^*™*1^ can be sampled.

#### 2.2.2 Combining sensor estimates

Each sensor has its own local proportional saturation *g*_*j*_(*x*), which gives the proportional water concentration relative to the value at minimum *x*, with shape determined by physical heterogeneity. Each sensor can be used as an independent estimate of water concentration as a proportion of maximum. At the same time, the total water content of the container is a global property of the system. Our method estimates, per sensor, the total water content. Given *N* sensors with voltage measurement vector **x**, each inverse sensor response is itself a random variable. We refine our single-sensor model as:

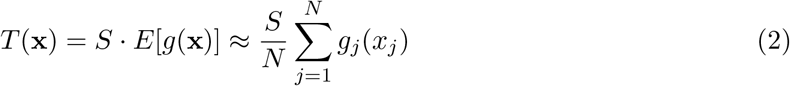

allowing *S*, a global quantity to be estimated jointly, with separate estimates for the local relative saturation functions *g* per-sensor.

#### 2.2.3 Model

The sensor response has two plateaus, one at dry (low SWC) and one at wet (high SWC). Different functions may perform better for different subsets of the domain. To model this, we consider two components: an exponential shape with a maximum of 1, which may differ between sensors, and be informed by both a prior normalized calibration curve and Δ*T*, and a joint scale, which is informed by Δ*T* . We use an exponential shape and do not attempt to model the dry end shape, instead using an intercept term. Given that the response function becomes flat at saturation, the inverse response function becomes vertical at full saturation. We do not attempt to extrapolate beyond the measured sensor minimum because small errors in assumed *X*_*min*_, which may vary between sensors, can result in large errors in estimated SWC where the curve is nearly vertical. We model the inverse mapping from sensor reading to water volume:

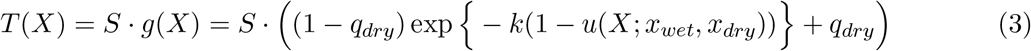

Where *u*(*X*_*t*_) is a linear rescaling of [*x*_*wet*_, *x*_*dry*_] to the interval [0, 1]. The shape intercept parameter *q*_*dry*_ allows the normalized curve *g*(*x*) to be squashed into the interval [*q*_*dry*_, 1.0]. The parameter *S* is equal to the total water volume at the minimum sensor measurement, and can be related to volumetric SWC as:

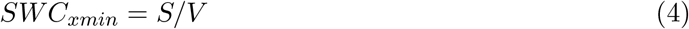

**Table 1:** Summary of Prior Distributions for Model Parameters.

| Parameter | Prior Distribution | Description |
| --- | --- | --- |
| $S$ (Scale) | Normal | Weakly informative; roughly estimated by user |
| $k$ (Exp. shape) | Improper | Unconstrained shape, unintuitive for user |
| $q_{dry}$ (Intercept) | Uniform(0, 0.5) | User-specified, default chosen from experiment |
| $\sigma^2$ (Variance) | Improper | Accommodates noise sources including error in water measurement |

#### 2.2.4 Bayesian estimation

To combine the chord data (*X*_*i,start*_, *X*_*i,end*_, Δ*T* ), dry anchor sample *X*_*i*_(0) = *h*^*™*1^(*T* = 0), prior assumptions on the parametric form of *h*^*™*1^, optional prior calibration points, and the unknown scale parameter *S*, we adopt a Bayesian framework. We assume lognormal dosage samples. For a single sensor, the response to dose Δ*T* is modelled as:

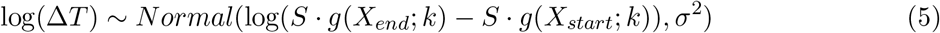

With the target parameters *θ* = (*S, k, σ*^2^, *q*_*dry*_).

Estimation is performed with the *emcee* [5] package in Python. Given posterior samples, volumetric SWC is estimated by Monte Carlo sampling, given the independence of *V* and calibration model parameters.

### 2.3 Test system

The described method was demonstrated by deployment of a system of two capacitive moisture sensors into a single pot with a lime tree, *Citrus × aurantiifolia*, and monitored over time. Regular watering was undertaken during routine care with doses between 100 ml and 300 ml. A prior curve from previous gravimetric SWC calibration of general purpose potting compost was used, comprising 15 values ranging from 885 mV to 1973 mV, averaged from 2 sensors.

### 2.4 Simulation

Evaluation was undertaken on a simulated dataset with two sensors. Briefly, given a known modified exponential response function, as shown in Equation (3) with scale *S* = 270, fractional intercepts *q*_*dry*,1_ = 0.05 and *q*_*dry*,2_ = 0.1, exponential shape *k*_1_ = 5.0 and *k*_2_ = 10.0. Each sensor was given its own minimum *X*_*min*_, corresponding to saturation: *X*_*min*,1_ = 3.0, *X*_*min*,2_ = 2.5. A single *X*_*max*_ = 8.5 value was utilised. Chord data (*N* = 64) was simulated with multiplicative unit-mean lognormal noise with variance of *σ* = 0.2. Uninformative end-point only prior calibration curves and a single anchor point at (*X*_*max*_, 0) were used. A *Beta*(1, 3) prior was applied to both *q*_*dry*_ shape intercept parameters. Inference was run with a burn-in of 400, with 1000 samples taken post-burnin. In a second simulation, for exploring identifiability, a run with only beta-distributed prior on *q*_*dry*_ was used, without anchor points.

## 3 Results

Figure 4 shows a corner plot for estimated parameters from simulated data with noise. At a noise variance *σ* = 0.2 and *N* = 64, true parameter values fell within 95% HPD intervals. Importantly, the scale factor giving total system water at *X*_*min*_ was within 5% of the true value (*Ŝ* = 284.6, *S* = 270). Appendix Figure 5 shows the noise magnitude as a function of water dose Δ*SWC*. Appendix Figure 6 shows parameter traces and mixing after burn-in. As demonstrated in Appendix Figure 7, without anchor points, identifiability of *q*_*dry*_ relies on the beta-distributed prior, and banana-shaped joint posteriors are observed, showing dependency between *S* and both *q*_*dry*_ parameters.

For the deployment into the Lime tree (*Citrus × aurantiifolia*) test system, Figure 1 gives the inferred SWC time series for each sensor along with the combined estimate. Figure 2 gives the inferred calibration curves. Both figures demonstrate consistency of combined estimates between both sensors, and smaller 95% HPD intervals for joint estimates over single-sensor estimates, obtained through the averaging scheme described. Figure 3 gives the corner plot showing the joint posterior distributions, showing no strong coupling between parameters. In contrast to simulated data, the true values for parameters are not known. Although gravimetric assessment of SWC for a potted lime tree would be destructive, future work should use gravimetry for validation in a range of real systems.

**Figure 1.**
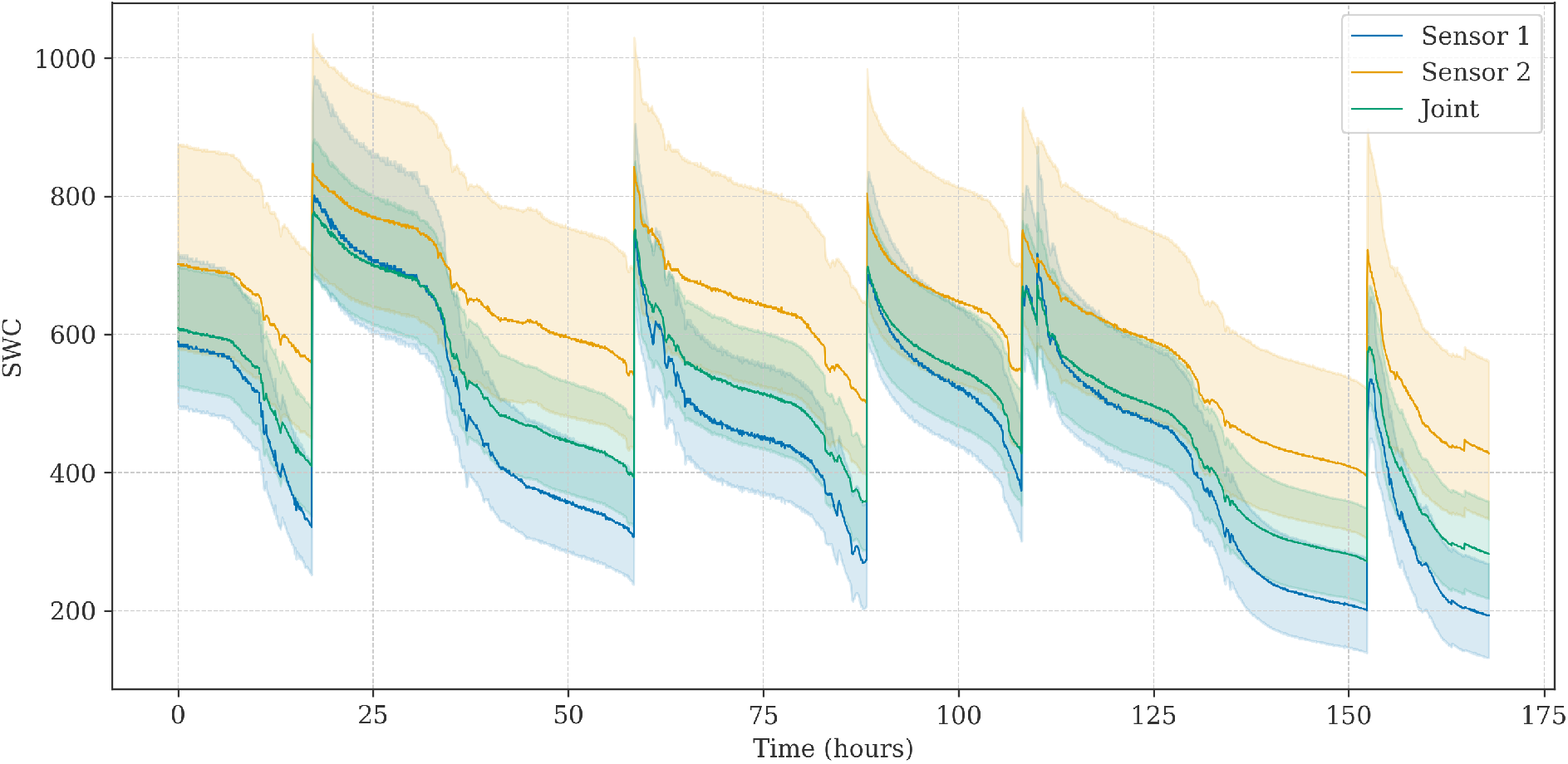
Estimated SWC over time from multi-sensor Bayesian calibration. Single-sensor estimation is shown with the combined sensor estimate. Shaded regions denote the 95% HPD.

**Figure 2.**
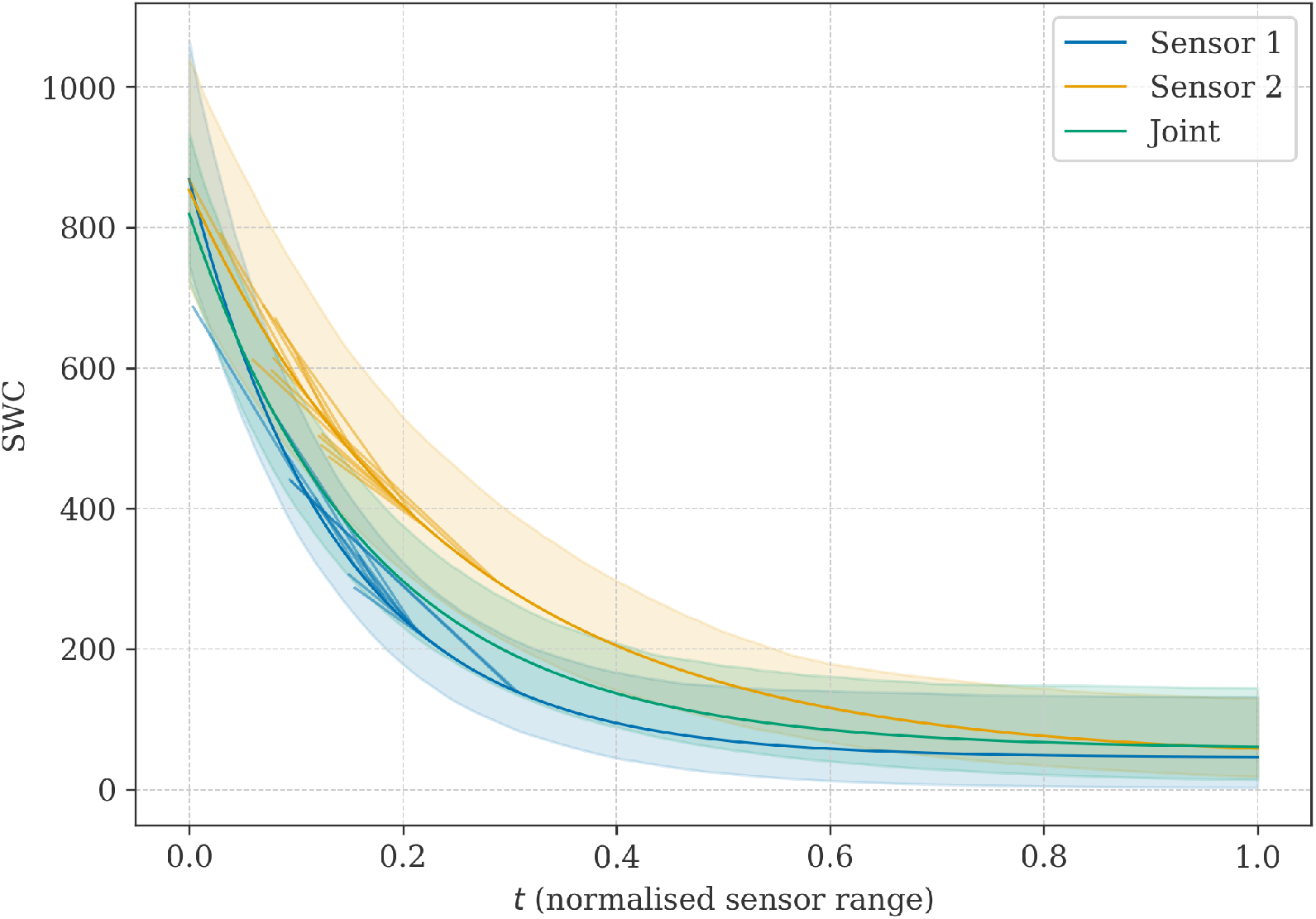
Calibration (response) curves per sensor and joint calibration, plotted against normalised sensor range *u ∈* [0, 1]. Chord segments show the observed watering events. Shading shows 95% credible intervals.

**Figure 3.**
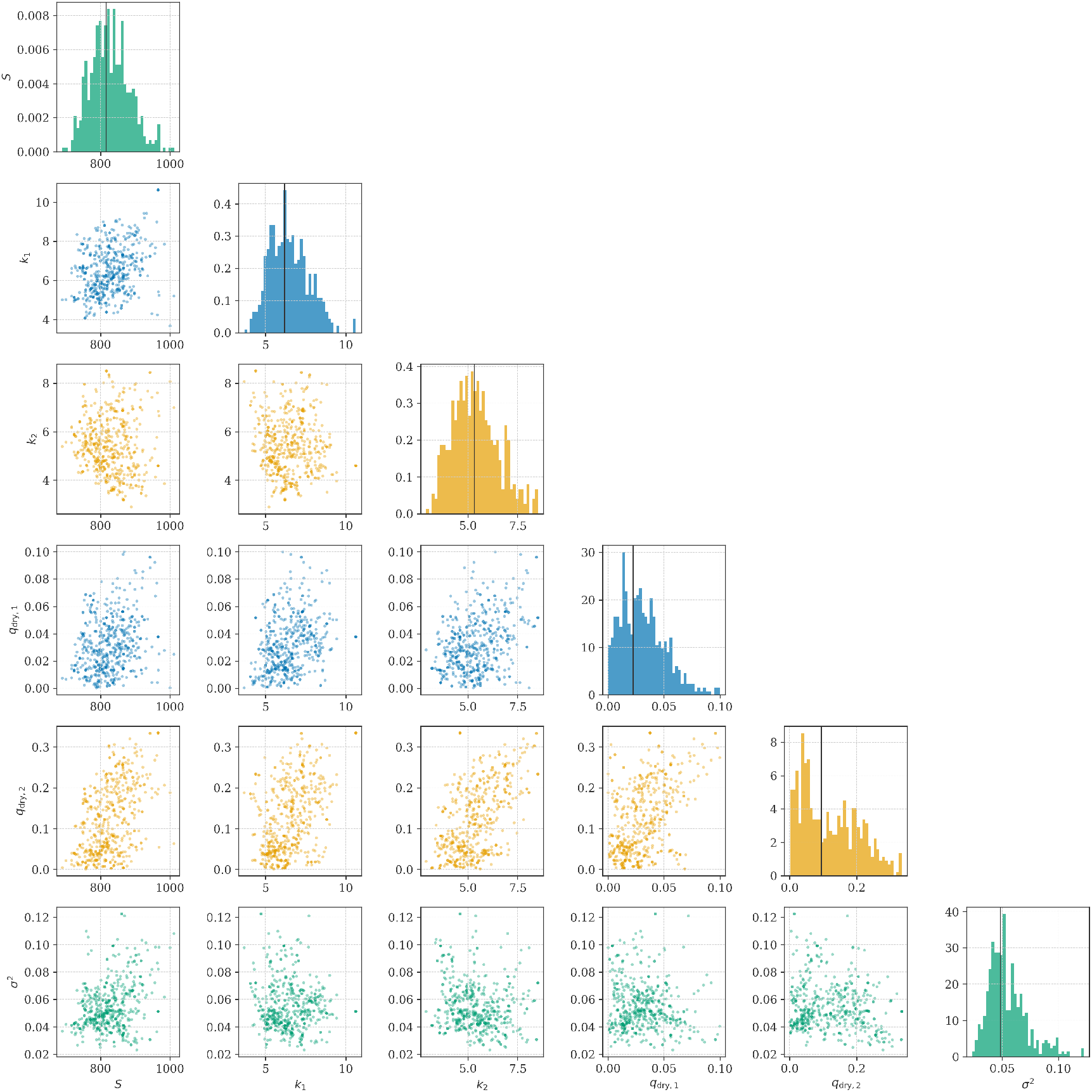
Corner plot from joint MCMC calibration with real data. Diagonal panels show marginal distributions for each parameter; lower triangle shows pairwise joint samples.

**Figure 4.**
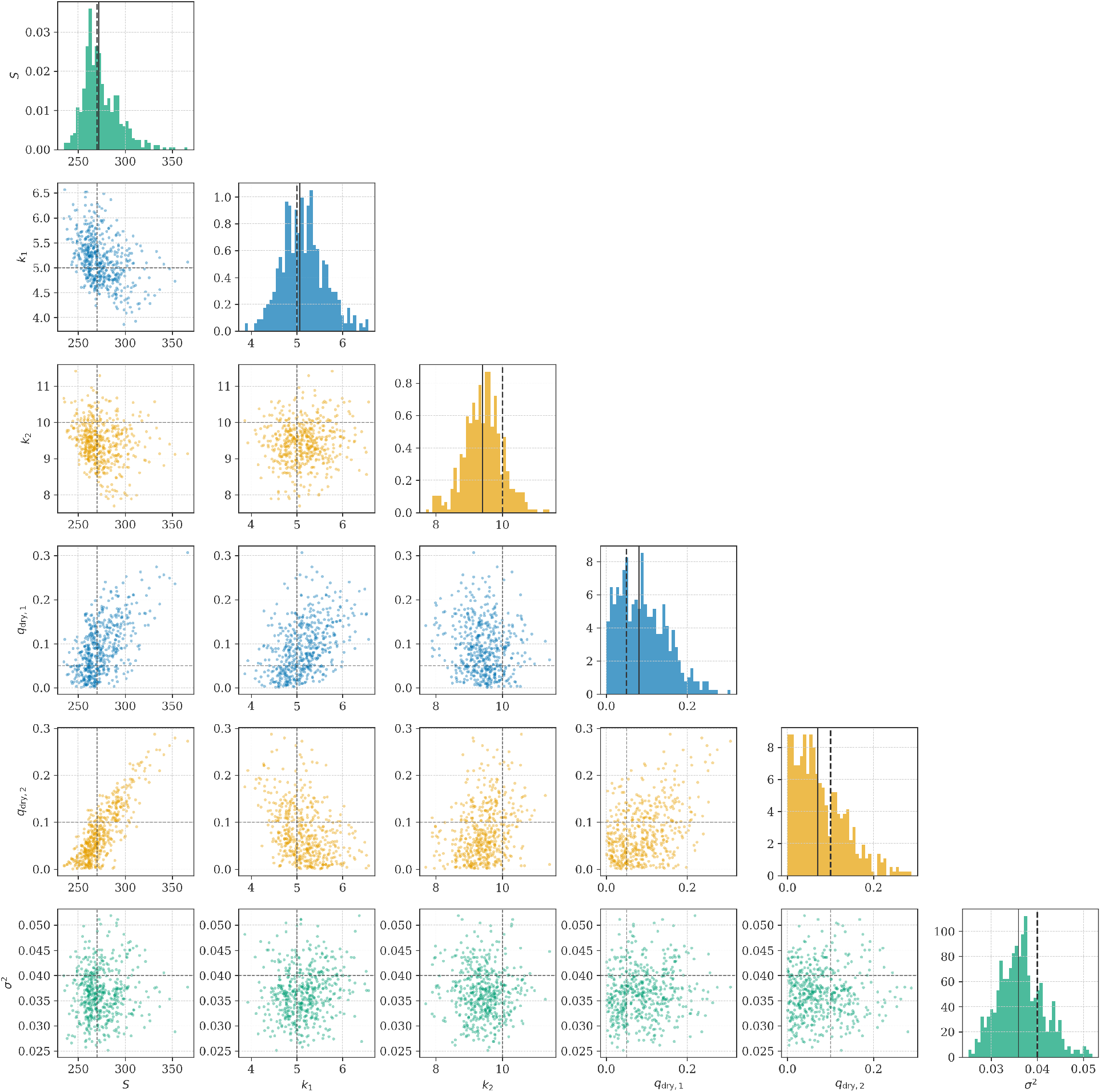
Corner plot from joint MCMC calibration with simulated data. Diagonal panels show marginal distributions for each parameter; lower triangle shows pairwise joint samples.

**Figure 5.**
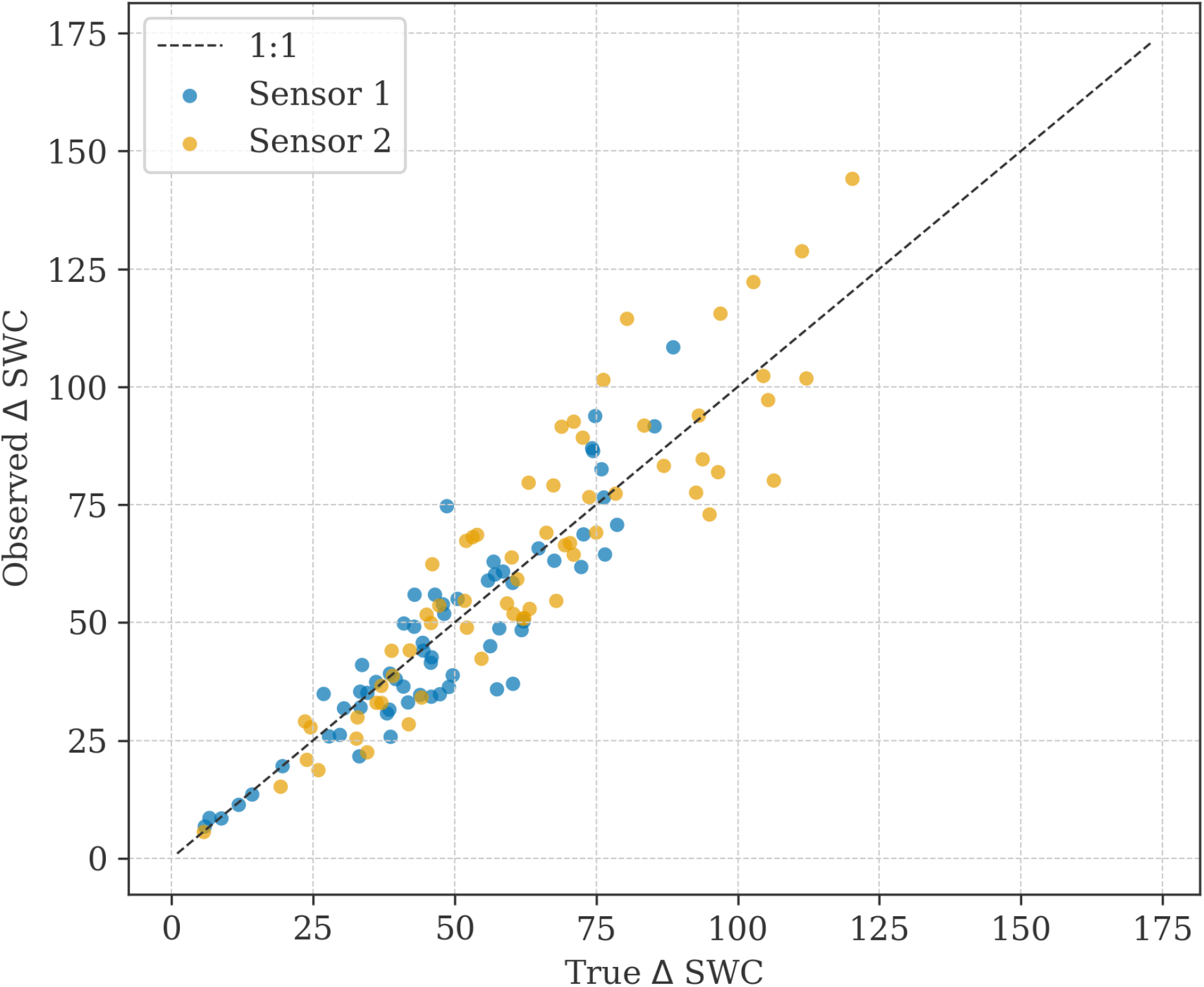
Observed change in soil water content with noise for simulated data versus true change. At *σ* = 0.2, multiplicative noise used during simulation mostly falls within 25% of the true value.

**Figure 6.**
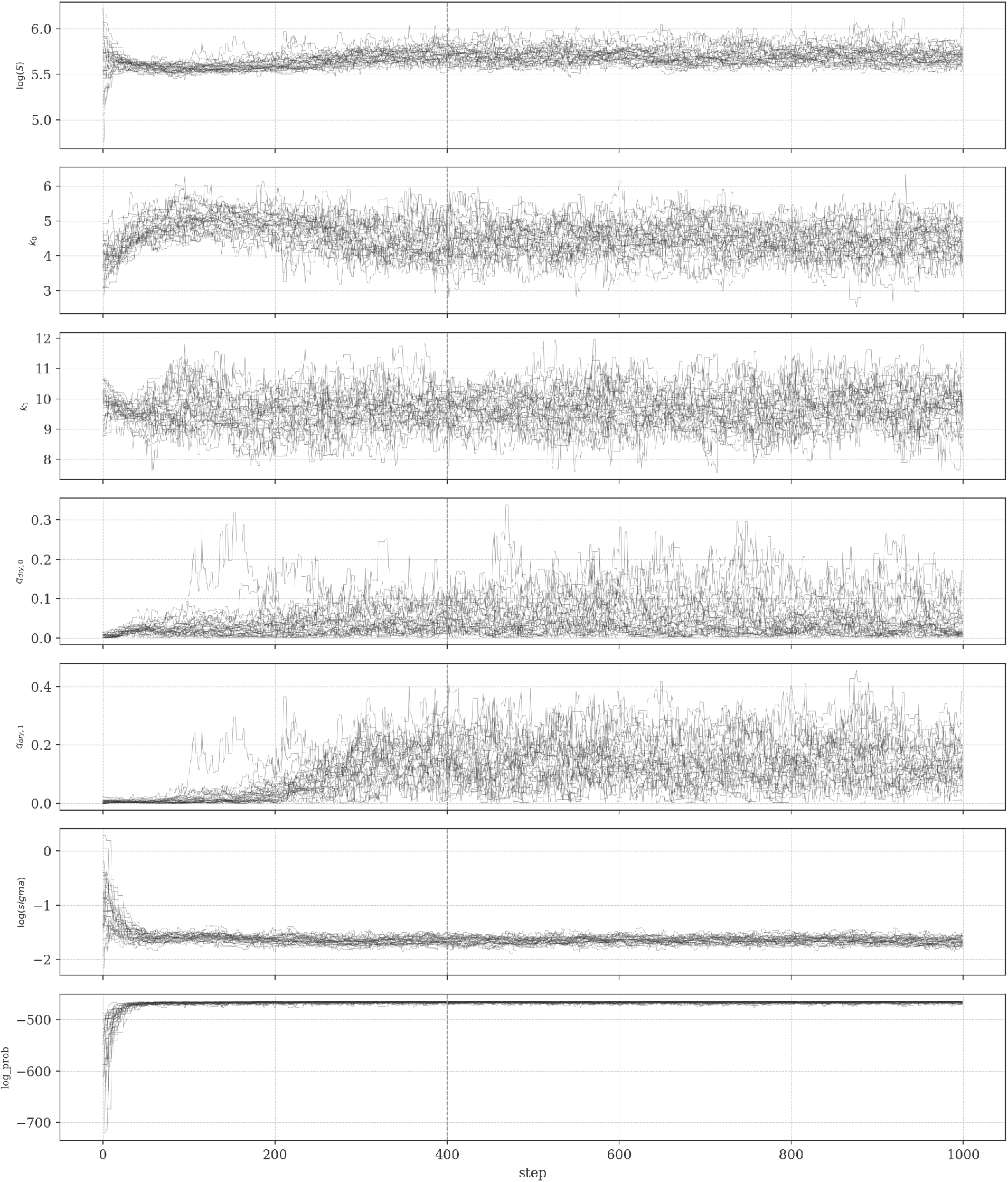
Traces for shared-sensor parameters during inference on simulated data, showing mixing after burn-in.

**Figure 7.**
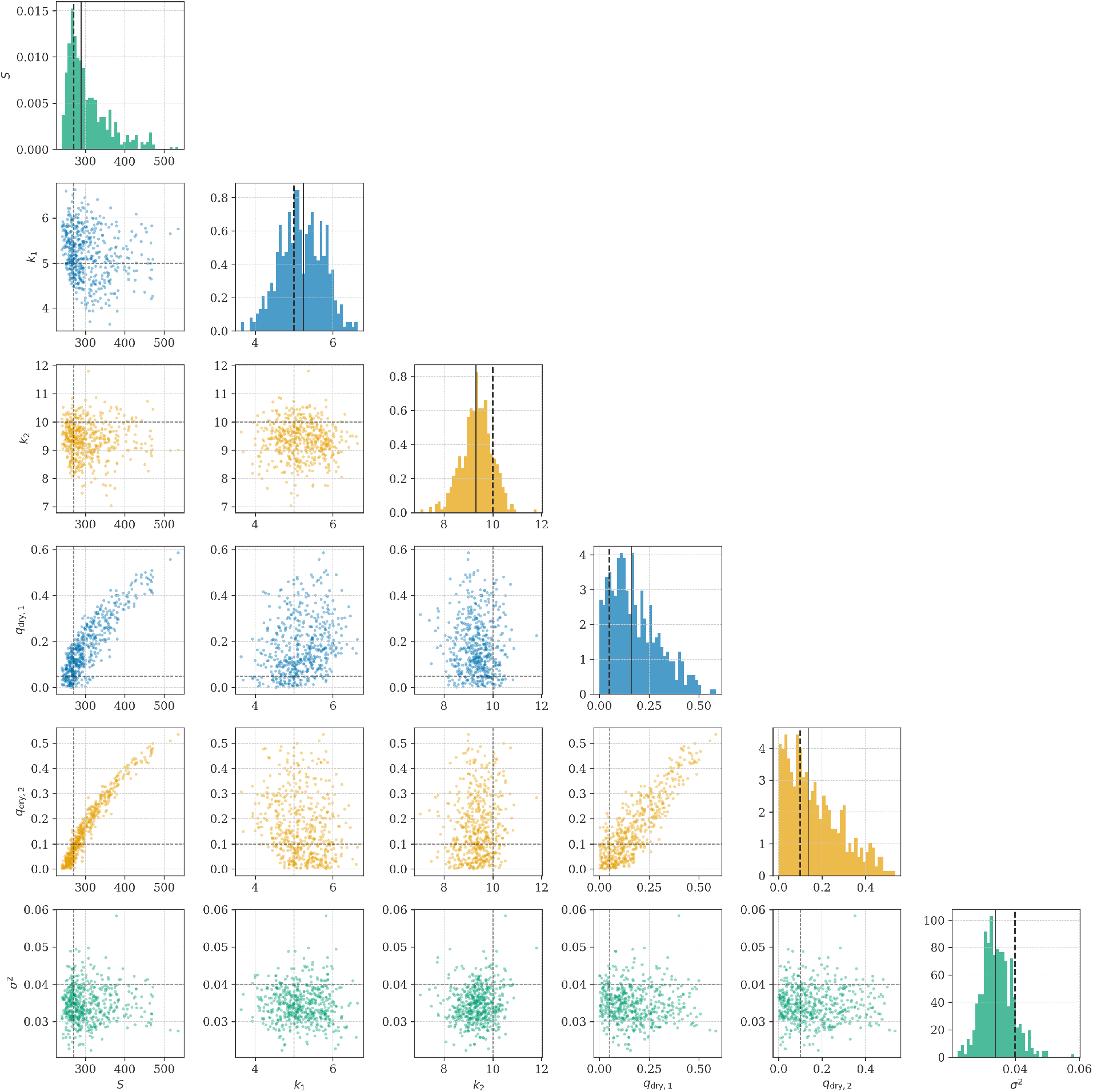
Corner plot from joint MCMC calibration with simulated data without anchor or calibration curve prior. In this case, with only a beta-distributed prior informs *q*_*dry*_, banana-shaped sample distribution in the joint posterior. Diagonal panels show marginal distributions for each parameter; lower triangle shows pairwise joint samples.

## 4 Discussion

The central concern for future work is to validate across real soil samples with a robust methodology such as gravimetry. In order to demonstrate the central utility of our method, future experiments must apply to a range of soil types and plant-tolerable SWC ranges, to demonstrate sufficient flexibility and alleviation of the practical difficulty of ESP32-based sensor systems. It is critical that methodology designed for calibration is tested for performance against real data with known SWC values. Although simulated data is presented, demonstrating reconstruction of known response functions, future work should apply the method to a range of real soil types and volumes to assess robustness in different systems.

The results demonstrated highlight a particular concern on identifiability. For the real data use-case, a prior curve was utilised using manual calibration data. For the simulated data cases, in the absence of a calibration curve prior, banana-shaped joint posterior distributions are structurally expected for *S* and *q*_*dry*_ for each sensor. Without prior information, chord data, as integral data, cannot inform the intercept, only scale and shape. There are a few resolutions to this. It is possible to utilise a prior, known anchor points (such as a single sensor measurement for dry soil), or to fix the intercept to zero. The simple approach of taking a zero-intercept is reasonable but cuts off the dry-end of the curve. This will have variable impact depending on the region of the curve that a given plant is kept in. A second identifiability problem arises when the subset of the domain covered by chords is low. If noise is high, then as the section of the domain of *X* covered by data shrinks, it is expected that it is harder to identify the shape parameter.

The advantages of our proposed methodology include: credible intervals on output SWC time series, without which point estimates are hard to assess; the ability to compute derived quantities such as velocity from posterior samples; joint estimation across sensors, so that a network of sensors can contribute to a central estimate; integration of prior information, such as from calibration on similar soil types; applicability to cheap ESP32 sensor prototypes.

## 5 LLM disclaimer

This paper was written without LLM generation. LLM generation was used for code and research assistance.

## 6 Appendix

